# The use of ATR-FTIR spectroscopy to characterise crude heparin samples by composition and structural features

**DOI:** 10.1101/744532

**Authors:** Anthony Devlin, Lucio Mauri, Marco Guerrini, Edwin A. Yates, Mark A. Skidmore

## Abstract

Production of the major anticoagulant drug, heparin, is a complex process that begins with the collection of crude material from a dispersed network of suppliers with poor traceability, an issue that was made apparent in 2007-2008, when batches of heparin were contaminated deliberately in the supply chain, resulting in over 100 deaths in the US alone. Several analytical techniques are used currently for the characterisation of pharmaceutical grade heparin, but few have been applied to its crude counterpart. One exception is NMR spectroscopy which was used to study crude heparin (2017), however, owing to the high set-up and running costs, as well as the need for skilled technical operators, the use of NMR at crude heparin production plants is unviable. An alternative, practical, spectroscopic method is attenuated total reflection Fourier transform infrared spectroscopy (ATR-FTIR) that is user-friendly, economical and, importantly, requires little specialised training or sample preparation. Using a top-down chemometric approach employing principal component analysis, ATR-FTIR spectroscopy was able to distinguish crude heparins based on their similarity to pharmaceutical heparin, as well as on their compositional and structural features, which included levels of sulphation, the extent of related conformational changes, as well as the quantities of chondroitin and dermatan sulphate present. This approach lends itself to automation and will enable users and regulators to undertake quality control of crude heparin during manufacture. The method requires only economical, portable equipment and little specialised training, bringing the high-quality analysis of crude heparin within reach of both manufacturers and regulators for the first time.

## Introduction

Heparin is a member of the glycosaminoglycan (GAG) family - a group of linear, polydisperse polysaccharides that exhibit numerous vital biological roles, many of which stem from the structural resemblance of heparin to the ubiquitously expressed mammalian cell-surface polysaccharide, heparan sulphate (HS)^1^. Nevertheless, heparin exhibits several activities, mediated particularly through interactions with proteins of the blood clotting cascade, including antithrombin III, factor IIa (thrombin) and factor Xa, which have led to its widespread use as a clinical anticoagulant and its inclusion on the World Health Organisation’s list of essential medicines ^2^. The sequence of heparin is characterised by a disaccharide repeat of the amino sugar alpha-D-glucosamine (A), which is predominantly N-sulphated (NS), but may be N-acetylated (NAc) and O-sulphated at position 6 (6S)or, more rarely, 3-O-sulphated) (3S) and a uronic acid residue (either beta D-glucuronic (G) or alpha L-iduronic acid (I)), which may be O-sulphated at position 2 (2S). The sulphation of disaccharide units may vary throughout the polysaccharide and a measure of the average sulphation of each disaccharide unit, dubbed degree of sulfation (DoS) is often used to compare samples, however, heparin disaccharides typically consist of 70-80 % I2S residues and GNS and 6S residues, mixed with lower levels of other constituents including D-glucuronate, unsulphated I (IOH), GNAc and some G3S, to generate considerable sequence diversity^3,4^,^5^. The structural diversity of pharmaceutical heparin is compounded further by variations in molecular weight, compositional differences between heparin from different organs, individual animals and different species^6^, and is amplified by the practice of amalgamating the material obtained from many individual animals before processing^7^.

Heparin for medical use is obtained primarily from porcine or bovine intestine, and can be sourced from ovine intestine and bovine lung as well, however, owing to bovine spongiform encephalopathy (BSE) in the late 1980’s and early 1990’s, predominantly heparin from porcine intestinal mucosa is used in Europe and North America^2^. Typically, the porcine intestine is collected and processed in slaughterhouses approved by regulatory authorities, where the mucosa is scraped from the intestines. The mucosa is then hydrolysed, and the resulting glycosaminoglycan mixture subsequently loaded onto an anion exchange resin before transportation to crude heparin processing facilities, where the loaded resin is washed with water and a solution of low-ionic strength. The enriched heparin fraction is then removed from resin with a high salt concentration and filtered, precipitated, and vacuum dried to form the substance formally known as crude heparin - a mixture, typically comprising approximately two parts heparin and one part “other” substances, including other GAGs such as chondroitin sulphate (CS), dermatan sulphate (DS), or heparan sulphate (HS). This mixture also contains nucleic acids from the parent species, residual proteins and a range of bound small molecules and ions. Additional processing is required to remove most of these contaminants^8^. The crude product then undergoes oxidative and alkaline treatments to remove residual colour, and to inactivate endotoxins and viruses that remain following extraction, resulting in the final pharmaceutical product^8^.

While the steps that create pharmaceutical quality heparin from crude heparin are performed under cGMP regulations, the traceability of the porcine intestine suppliers that create the crude heparin is poor and, since many sites are involved, the risk of contamination from an unknown source is greatly increased^7^. This is particularly pertinent in modern heparin manufacture, following the contamination of pharmaceutical heparin in 2007-8, dubbed the “heparin crisis”, which was caused by the introduction of material, much of which was an unnatural, chemically modified chondroitin sulphate (over-sulfated chondroitin sulfate; OSCS) into the supply chain, and resulting in approximately 150 deaths and 350 other adverse events in the USA alone ^8,9^. Subsequently, new tests were introduced, including rudimentary proton NMR spectroscopy, and HPLC, to the heparin monograph in an attempt to reduce the risk of product adulteration^10^,^11^. Nevertheless, significant quantities of alien material can persist.

Many studies have been undertaken with the aim of detecting contamination (almost always with OSCS^12–16^, even though a very wide range of other potential future contaminants, whether accidentally or deliberately introduced, can be envisaged^17^) in pharmaceutical heparin samples, without first defining what level of structural variation is observed between *bona fide* heparin samples. In the case of pharmaceutical heparin, this issue was first identified and addressed in 2011 using proton NMR^18,19^ and, in 2017 the first steps in this direction were taken for crude heparin using NMR spectroscopy and strong anion exchange HPLC (SAX-HPLC), coupled with principal component analysis (PCA)^7^. Using this approach, it was possible to characterise crude samples by their composition and structural features. Recently, the cheaper and practically more straightforward spectroscopic method, ATR-FTIR, combined with PCA, was shown capable of distinguishing various GAGs, heparins contaminated with OSCS and other carbohydrates from heparin species^15^.

Here, 73 of the original 75 crude heparin samples (taken from those used in the 2017 study^7^) were analysed with ATR-FTIR, and the viability of this method for use in the characterisation of crude heparin was examined. This approach is particularly pertinent because, when the dispersed nature of the heparin supply chain is considered, the simple machine set-up, straightforward sample preparation, short acquisition time and relatively undemanding level of training or expertise required (compared with NMR and SAX-HPLC) makes ATR-FTIR a strong candidate for practical use by manufacturer and regulator alike.

## Results

### (i) PCA analysis of the ATR-FTIR spectra reveals the similarities and differences between crude heparins and other GAGs

The crude heparin samples were compared to the GAG library and heparin libraries established in ^15^ using PCA to help observe their inherent similarities and differences (**Fig. 1**). The crude heparin samples appeared distinct to the *bona fide* pharmaceutical heparins, but remain close to them. The most heparin-like samples (dark red) are found well inside the region which correlates with heparins in **Fig.1A**. When compared solely to the heparin library, the crude heparins appear initially distinct in PCs 1 and 2; the most heparin-like samples again appear in the region correlated with heparins. In components 3 and 4 (**Fig. 1Bi**), the separation seen between heparin species in previous work^15^ is retained; the crude heparins covering the PMH region of the PMH, BLH and OMH cluster, now appearing in the gap between the BMH and PMH-OMH-BLH clusters (**Fig.1B**). ATR-FTIR with PCA can therefore distinguish crude heparins from other GAGs, and pharmaceutical heparin.

**Figure 1.**
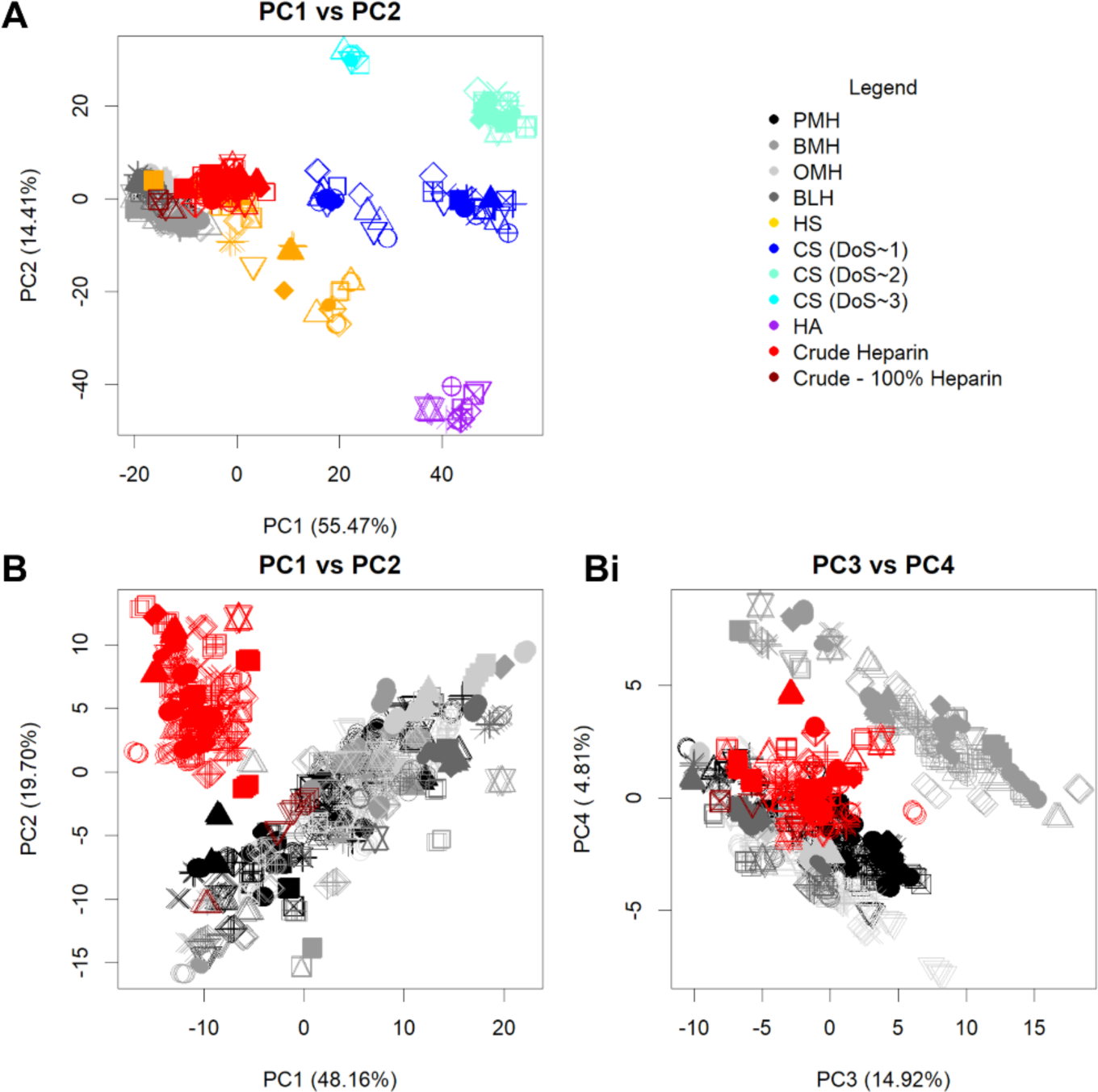
PCA analysis of the ATR-FTIR spectra reveals both the differences and similarities between the crude heparins, bona fide heparins and other GAGs. **A** PCA analysis of 73 ATR-FTIR spectra of different crude samples (red, dark red for crude samples that most closely resemble pharmaceutical heparin), 185 heparins comprising: 69 PMHs (black), 55 BMHs (grey), 33 OMHs (light grey), 19 BLHs (medium grey), 31 HSs (gold), 44 chondroitin (including 13 CSA samples, 16 CSC samples and 21 DS (blue for monosulphated chondroitin samples and teal for disulphated chondroitin samples), 11 HA (purple) and 6 OSCS (aquamarine) samples. **B** PCA analysis of the 73 crude heparins and 185 heparinsusing PCs 1 and 2. **C** PCA analysis of the 73 crude heparins and 185 heparins using PCs 3 and 4.

### (ii) Assigning ATR-FTIR bands from the literature

Before examination of the ability of ATR-FTIR spectroscopy to distinguish structural features of the crude library established in ^7^) rudimentary band assignments for ATR-FTIR of heparin were obtained. There is currently no completely assigned ATR-FTIR spectrum of heparin because of the complex and overlapping bands that characterise the ATR-FTIR spectra of heparin samples. This complexity arises from the natural heterogeneity and complexity of the material, but several groups have made efforts to assign some of the bands (**Table 1**). Assigning IR bands is fundamentally challenging, as bands can involve numerous coupled modes and overtones, and the continuous nature of IR spectra mean that multiple bands overlap. Nevertheless, assignments have been made below 1000 cm^-1^. Here, features unique to the backbone of the molecule, such as the directionality of C-H bonds, 2S and 6S have been assigned to bands at 875, 800 and 875 cm^-1^ respectively ^21^,^22^,^27^. Some assignments have also been made above 1000cm^-1^ in heparin for S=O stretches and, symmetric and asymmetric carbonyl stretching at 1230, 1430 and 1635 cm^-1^ respectively^21,37^,^26,28^. Typically, one or two bands unique to the studied GAG are used for identification of the molecule^28^,^33^,^34^,^35^ but this is of little use for quantitative structural feature assignment. Many more assignments have been made for carbohydrates in general, and those -relevant to heparin are displayed in **Table 1**. Peaks unascribed to heparin can be assigned using these carbohydrate bands, and have been assigned in a similar fashion as they have been to other GAGs in^33^,^39^,^40^. It is usually assumed that the heparin and crude heparin samples can only contain certain bond types or functional groups (for example, stretches between 1600 and 1700 cm^-1^ can be assigned to C=C bonds or amine related bonds^20^, but unmodified heparin does not contain a C=C bond, therefore amine bonds are assigned) and it follows a pattern similar to other GAG IR spectra, however, some bands remain unassigned.

**Table 1:** IR band assignments for heparin from the literature.

| *Band Peak/Region /*<br>*cm<sup>-1</sup>* | *Structural feature/bond correlated with* | *References* |
| --- | --- | --- |
| 720 | Bound 2nd amine, N-H deformations | 20 |
| 800 | 2S | 21, 22 |
| 827 | 6S | 23,24 |
| 836 | C1-H | 25 |
| 850-950 | Deformation of C6 | 22,24,26 |
| 875 | Equatorial vs axial C-H | 27 |
| 890 | Coupling of C-O-S and ring C-O-C | 23,28 |
| 911 | C1-H | 25 |
| 913 | O-H | 25 |
| 930-1000 | Glycosidic bond, C-O-S stretches | 23,25 |
| 950-1170 | Stretching indicative of a pyranose ring | 29,30 |
| 1000-1200 | Characteristic carbohydrate C-O stretching and C-O-H bending modes | 31 |
| 1020-1050 | S=O symmetric stretching | 20,28 |
| 1030-80 | O-H bands | 27 |
| 1070-1150 | C-O-C asymmetric stretching | 20 |
| 1100 | Sulphate | 20,27 |
| 1180-1200 | N-sulphation | 28,32 |
| 1230 | S=O<br>Hp: 1245, 1222<sup>28</sup><br>CSC: 1240, 1220<br>CSA: 1240, 1230<br>DS: 1240, 1225 | 20,23,27,28<br>33, 34, 35 |
| 1230-1235 | N-sulphonate (Obscured by high OS) | 21,32,36 |
| 1240 | C=O stretching of NAc, and S=O Asymmetric stretching | 27,28 |
| 1210-1320 | C-O carboxylic acid | 20 |
| 1250-1310 | S=O | 22 |
| ~1340 | C-H bending | 20,25 |
| 1349 | O-H | 25 |
| 1382-89 | C-H stretch NAc | 20,21 |
| 1417-33 | Carboxylate stretching | 20,21,37 |
| 1429-32 | Carboxyl – NS interaction | 21,37 |
| ~1468 | C-H scissoring (alkane) | 20 |
| 1530-1565 | Amide II vibrations (Including free amine) | 20,21,26,31,38 |
| 1540-1610 | C-O asymmetrical stretch carboxylate | 20 |
| 1550-(1650) | NAc deformation | 20,27,31,38 |
| 1718 | Acyclic carboxylate | 20 |
| 2800-3049 | Stretching of C-O/C-C | 30 |
| 3100-3700 | Bound OH | 20,27,30 |
| 3500-3700 | Free OH | 20,27,30 |

### (iii) Simple visual inspection of raw ATR-FTIR spectra cannot distinguish reliably between different features

The FTIR spectra, following baseline correction and normalisation (0-1) were compared visually against one another, particularly focusing on samples with the highest and lowest levels of different structural features in order to ascertain whether the FTIR spectra would change in a predictable or obvious manner (**Fig. 2**). The spectra of samples with high CS, high DS and both high CS and DS were compared against a randomly selected heparin sample from the heparin library (i.e a sample with 0% CS and DS content). There are noticeable differences between these spectra, particularly in the intensities of the peaks at ∼1450 cm^-1^ and 1600 cm^-1^ and an extra shoulder appears at ∼1430 cm^-1^. The split peak at ∼1000 cm-1 has a more accentuated split in the secondary maxima at ∼1050 cm^-1^. The intensity at ∼1150 is lower in these samples. The variable region after 3000 cm^-1^ is unpredictable for all samples due in part to free hydroxyl groups from environmental water. At the second maxima (∼1050 cm^-1^), the sample containing both high CS and DS displays, paradoxically an intensity between those of the samples with high CS and high DS levels (**Fig. 2A)**. The samples were compared in a similar manner for sulphation type, with the overall degree of sulphation (DoS), levels of NS vs NAc and levels of I2OS vs I2OH being compared in **Fig.2B,C**. These spectra display a high level of similarity, with differences in the split peak maxima of ∼1050 cm^-1^ correlating to neither high nor low levels of these features. The bands do not alter in an obvious manner within the raw spectrum for the structural features, due in part to the samples natural heterogeneity, compositions and the broad and complex bands of the ATR-FTIR spectra.

**Figure 2.**
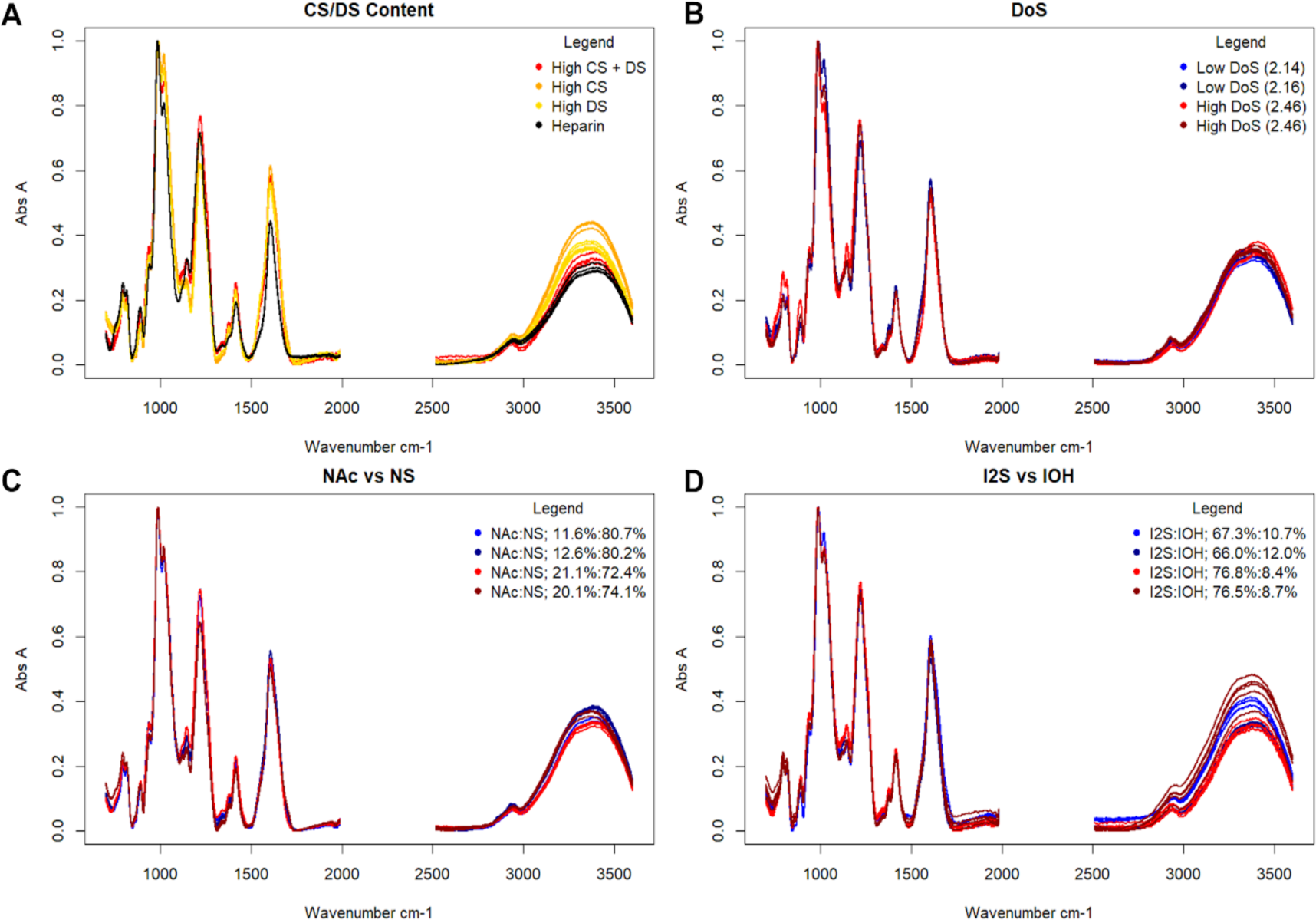
Illustrative examples of FTIR spectra of crude heparin samples containing relatively high and relatively low quantities of structural features within the studied library. **A** Comparison of sample composition. A randomly selected bona fide heparin (black) is compared with a crude heparin possessing high CS levels (red) DS levels (yellow) and both CS and DS levels (red). **B** Two samples with relatively low DoS (red) and two samples with two relatively low DoS (blue) are compared. **C** The level of NS vs NAc is compared, for two samples with relatively high NAc (red) and two with relatively low NAc (blue) and vice versa for NS level. **D** The level of I2S vs I2OH is compared, for two samples with relatively high I2OH (red) and two with relatively low I2OH (blue) and vice versa for I2OS level.

### (iv) Samples of crude heparin can be distinguished in terms of composition using components 1, 2, 3 and 5

The ATR-FTIR spectra of the 73 crude samples were subjected to PCA analysis in an effort to examine and distinguish the composition and structural features of the samples in a manageable format. The relative levels of 29 different structural features, established previously in ^7^ were superimposed onto the score plots whereby the quantity present is assigned by colour; low relative amounts beginning with **blue**, and high relative amounts ending at **green**. In cases where none of the structural feature was found (and the change from 0% to the next lowest percentage was a large step) **black** was used to indicate this.

Components 2 and 3 can be used to distinguish between the composition of each sample **(Fig. 3)**. Levels of DS and DS2S from 0-21% and 0-15% are separated reliably, across PCs 2 and 3. Components 1 and 5 are required to separate CS and CS6S from 0-8% and 0-50% respectively, due to the strong correlation of component 1 with 6S **(next section)**.

**Figure 3.**
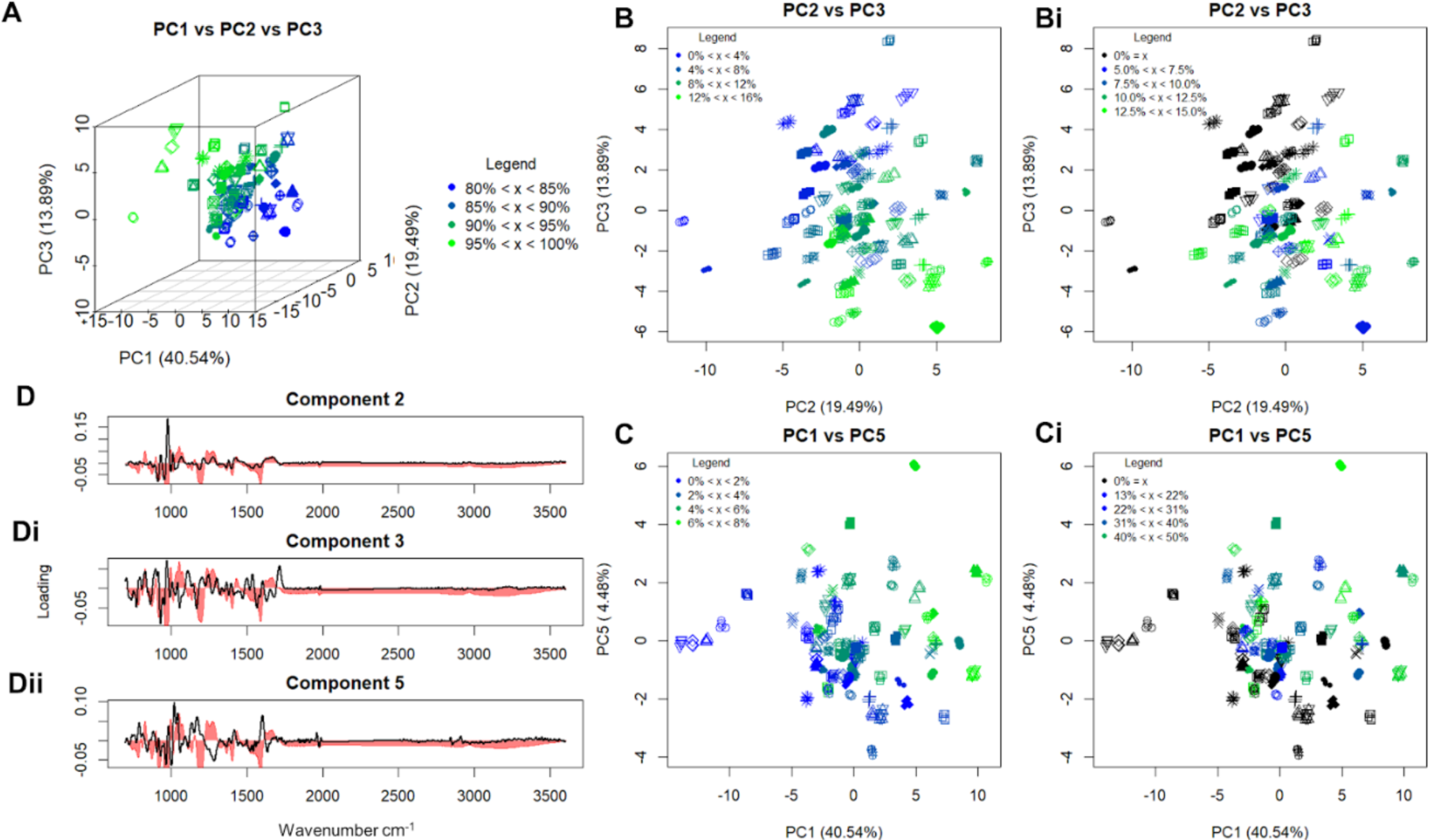
PCA analysis of the 73 crude heparin ATR-FTIR spectra correlate levels of other GAGs in the sample, which have been established previously^7^. **A** 3D score plot of PCs 1, 2 and 3, demonstrating the level of overall level of heparin present in the sample, from 80% (blue) to 100% (green) **B** Score plots of PCs 2 and 3 showing the levels of DS in the samples from 0% (blue) to 16% (green), **Bi**, Score plots of PCs 2 and 3 showing the levels of DS2S in the samples for 0% (black) and from 5% (blue) to 15% (green), **C** Score plots of PCs 1 and 5 showing the levels of CS in the samples from 0% (blue) to 8% (green), **C** Score plots of PCs 1 and 5 showing the levels of CS6S in the samples at 0% (black) and from 13% (blue) to 50% (green), **D** loading plots for components 2, 3 and 5. The loading (black) is overlaid over an average of the spectra of all 73 samples (red) showing those parts of the D2 spectrum that are responsible for the separations of DS and CS.

### (v) Samples of crude heparin can be distinguished by their structural features using PCs 1, 4, 5 and 6

The levels of different structural features were then compared and PCs 1 and 3 to 6 were found to correlate with various features. PCs 1, 3 and 5 correlated strongly with levels of sulphation found in the sample (**Fig. 4A**). Specifically, PC1 correlated to 6S levels, in both CS and heparin (**Fig. 3Ci, Fig 4C**) and PC 5 correlated strongly with NS/NAc levels (**Fig.4B**). Assignment of PCs 3 and 4 was more difficult, as they do not correlate with conventionally defined features. PC3 strongly correlated with both DoS and the % of polysaccharides possessing a linkage region, suggesting that it is relate to the overall structure of the molecules and the influence that interactions with each other have on the spectra (**Figs. 4D and Fi**) while PC4 correlated with the interactions of sulphates across glycosidic linkages, showing correlations in levels of ANS6X-I2S and G-ANAc6X (i.e the presence of two sulphates across the bond or not respectively). Interestingly, in examples containing a sulphate group, but no pairs of sulphates, such as ANS6X-G, there were no correlations found whatsoever across any components. PC6 separates by the presence of a functional group on C_2_ of the pyranose ring of either residue in the disaccharide unit, either as a −2S or an -Nx, there is a particularly strong separation for levels of ANS6X-I2S due the presence of both of these features at once. PC6 also correlated with NS/NAc levels, but another component is required for meaningful separation (data not shown). Structural features, when compared with their counter features (for example NS vs NAc) separate orthogonally across these components as would be expected. ATR-FTIR and PCA can therefore be used to distinguish crude sample composition, and the structural features of constituents of the sample, including, in some cases, the levels of sulphation of three different molecules in the sample.

**Figure 4.**
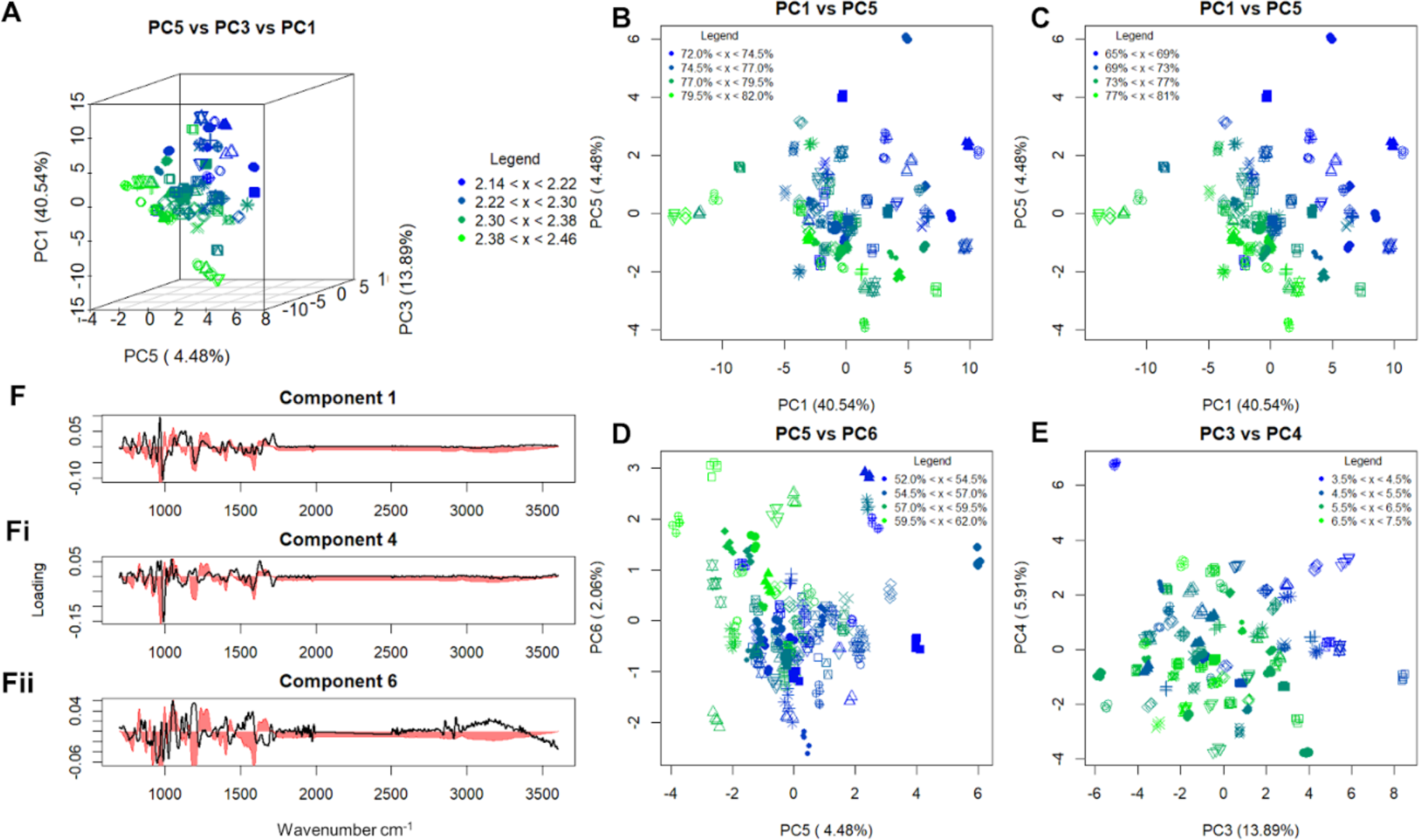
PCA analysis of the 73 crude heparin ATR-FTIR spectra allows correlation of distinct features, the amounts of which were established in ^7^. **A** 3D score plot of PCs 1, 3 and 5, demonstrating the average DoS in each sample, from 2.14 (blue) to 2.46 (green) **B** Score plot of PCs 1 and 5, demonstrating the levels of ANS6X and subsequently ANAc6X in the sample, 72%/21% (blue) to 81%/11% (green) for the level of ANS6X/ANAc6X respectively. **C** Score plot of PCs 1 and 5, demonstrating the levels of A6S in the sample from 65% (blue) to 81% (green). **D** Score plot for PCs 5 and 6 demonstrating the levels of ANS6X-I2S in the sample from 52% (blue) to 62% (green). **E** Score plot of PCs 3 and 4, demonstrating the levels of saccharide chains that still retain their linkage region from 3.5% (blue) to 7.5% (green) **F** loading plots for components 1 and 5, showing which parts of the spectrum are responsible for these separations.

The loading plots for each of these components, which show the regions of the original spectrum that are being examined in their respective components, were used in tandem with an average spectrum of all 73 samples to visualise those regions of the spectra responsible for separation **(Fig.3D and 4F)**. PC1 correlates with −6S, its loading plot contains positive values at 800 and 890 cm^-1^, corresponding to −6S levels and C-O-S, C-O-C interactions respectively, both a strong positive and negative spike at ∼980 cm^-1^ and a strong positive spike at 1100 cm^-1^, both associated with sulphation in heparins. PC2 has a strong positive spike at ∼1000 cm^-1^ in the area of the spectrum that differs the most between high/low CS/DS levels in crude heparins (**Fig. 2A**). The loading plot of PC3 is dispersed across the spectrum, as are most of the sulphate stretches. The loading plot of PC4 contains both a negative spike at ∼1000 cm^-1^ and an accented region at ∼1600 cm^-1^, associated with -OS and -NS respectively, correlating with the presence of both features or neither. In PC5 which is responsible for levels of NS/NAc, there is a positive cluster at ∼1200 cm^-1^ and ∼1500 cm^-1^, which are associated NS and, Amide and NAc vibrations respectively. PC6 correlated with the presence of a feature at C2 of the pyranose ring and had a dispersed loading plot, with its largest regions being found at 1100 cm^-1^ and 1550 cm^-1^ which are responsible for sulphate groups and amide II groups respectively (both of which can be linked to C_2_ functional groups) **(Figs. 3D and 4F).** ATR-FTIR with PCA can be used to highlight regions responsible for structural variation with good accuracy. It also shows that small structural changes have effects across the spectrum, explaining why it is difficult to distinguish the spectra without multivariate analysis. This means that analysts can use loading plots to identify regions associated with specific functional groups or interactions. Furthermore, the band assignments made in **Table 1** agree with the ATR-FTIR loading plots, when combined with information from NMR/SAX-HPLC analysis^7^, allowing greater confidence in the band assignments and the results shown here.

## Discussion

Ultimately, the aim of this work was to explore whether ATR-FTIR spectroscopy can be used to define crude heparins and this was achieved. One of the major issues facing crude screening the lack of any real regulation for it, meaning that some crude samples may in fact be a blend of very raw material coupled with pharmaceutical grade-heparin^7^. The authentic crude heparins studied here were shown to have their own place in the GAG library and to possess unique spectral features; the most heparin-like appearing in the heparin region and displacement of samples in the group depending on their composition, while also remaining separate and distinct from all other compounds. Any crude sample can be input into this analysis, and its similarity to heparin and other, recognised crude samples, could be identified, enabling quick and effective quality control and sample screening at crude heparin manufacturing sites, and allows users to begin to define what is a real crude heparin and what is a blend of other products (**Fig. 1**).

At first glance, ATR-FTIR did not appear to be of use for structurally defining crude heparin populations (**Fig. 2**). This was further compounded by the lack of full IR spectral assignment of heparin and other GAG molecules, as band differences could not be assigned to specific features or functional groups. With multivariate analysis of these spectra, the subtle differences in band thickness, intensity and location could be distinguished in a meaningful manner. By utilising a top down approach with PC loading plots and adding a third dimension with respect to structural or compositional features, one can distinguish sample composition such as CS and DS levels and percentage of structural features such as %NS, %OS (of heparin, CS and DS in the mixture) as well as even more complex interactions between functional groups. The complexity of the loading plots highlights the depth of the spectral bands and their relations to one another, PCs 1 and 5 for example separate A6S and ANS levels respectively, but the best separation for both features is found using both their respective components - a fact that may be explained in41, where it was shown that ANS is commonly also 6-sulphated in porcine samples, hence the best separation is found when both features are separated. This suggests that through combinations of components, the analyst may be able to distinguish levels of different saccharides, for example, the combination of PCs 1 and 5 may distinguish ANS6S levels. Further exploration of this is needed for validation however.

Later components seem to separate by more complex relationships. PC 6 separates by the presence of functional groups on both C2’s of a disaccharide unit or neither, but no separation is found for just one functional group per disaccharide. It suggests that the separation is correlated by ring or bond geometries, which are influenced by these functional groups. PCs 7, 8 and 9 offered no significant separation for features that were defined with NMR and SAX-HPLC, but the repeats of samples in the analysis remained somewhat close to each other, suggesting that there is still meaningful information to be found in these components. For example, levels of epoxide formation are poorly recorded with these techniques, hence samples cannot be grouped by this feature, but they may be separated by these components^41^. Repeats of the same sample in PCs 10 and above begin to separate distinctly, creating large intra-sample differences, hence these components show no meaningful separation.

Importantly, band assignments are not needed *a priori*, so long as the samples are grouped in a meaningful way by the analyst. However, once rudimentary band assignments were established, they agreed with the outputs produced. This still does not currently allow for assignment of unknown bands, as typically more than one component is needed for good separation of features, some of these components overlap for different defined features, it is unclear as to whether the separations are due to the structural features, or their effects on other bonds/regions and the analyst here is limited by what NMR and SAX-HPLC can and cannot determine, hence why PCs 8 and 9 were not assigned here.

In conclusion, ATR-FTIR can be used to define crude heparins based on “normal” crude heparins, and it may also be used to establish the structural features of the analysed compounds based on a libraries constituents. The multivariate approach provides a means for rudimentary band assignment in heparin IR spectra, however, a more systematic methodology is needed for true elucidation. Regardless, a rudimentary band table for heparin IR spectra from the current literature is also provided. Owing to the ability of ATR-FTIR spectroscopy to determine structural features, discriminate between heparin species^15^ and to be used in the field with fast set-up and spectral acquisition, the approach could prove fruitful for crude manufacturing quality control and help to prevent further crises.

## Methods

### Polysaccharide samples

Crude polysaccharide samples were gifted by the FDA. Other library polysaccharides were obtained as in ^15^.

### NMR/SAX-HPLX

Full details of these can be found in ^7^

### FTIR sample preparation

Prior to spectral acquisition a small amount of sample prep must be performed. 1-10 mg of dry sample was taken and 1 ml of deionised water added. The solution was frozen at −80°C and lyophilised overnight. It is important that the same drying technique is used for all samples to minimise variation from this source. Care was also taken to use the same type of tubes, to reduce any potential erroneous spectral features that arise from different drying rates associated with tube shape/volume.

### ATR-FTIR

Samples were recorded using a Bruker Alpha I spectrometer in the region of 4000 to 400 cm^-1^, for 32 scans at a resolution of 2 cm^-1^ (approx 70 seconds acquisition time), 5 times. A background spectrum was taken prior to each sample, using the same settings as for sample acquisition but with a completely clean stage. 1-10 mg of dried sample was placed upon the crystal stage of the instrument, ensuring that the entirety of the crystal was covered. Sufficient sample was employed to ensure that at least 5 µm of crushed thickness was obtained, as this is the extent to which the effervescent ATR wave penetrates. The instrument stage was cleaned with acetone and dried between samples. Spectra were recorded using OPUS software (Bruker) and exported in CSV format.

### FTIR processing

All processing and subsequent analysis was performed on an Asus Vivobook Pro (M580VD-EB76), using an intel core i7-7700HQ. Spectra were imported into R studio v1.1.463 prior to a preliminary smooth, employing a Savitzky-Golay smoothing algorithm (*signal* package, ***sgolayfilter***), with a 21 neighbour, 2^nd^ degree polynomial smooth, facilitating the improved accuracy of later corrections.

### Baseline correction

While a background spectrum is taken prior to each new sample, perturbations in the air can still occur during spectral acquisition. To further reduce the effects of these perturbations, each individual smoothed spectra underwent a custom baseline correction using a 7^th^ order polynomial. First, the spectra were separated into 6 equidistant buckets, with the minimum absorbance value for each of these buckets and their relevant x axis values taken. The start and end values for the spectrum were added to these values and, from the resultant 8 x-y pairs, the coefficients for a 7^th^ order polynomial were generated using *base* R ***lm*** function. The baseline was formed utilising the generated coefficients from the original x-axis and the calculated baseline subtracted from the smoothed spectrum. To remove the effects of the variable quantity of sample used to record a spectrum (which cannot be regulated, because there is no control over the amount of sample placed into contact with the ATR crystal), the corrected spectra were normalised (0-1) using the equation:

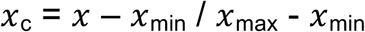

Where x is the value to be corrected, xc is the resultant corrected value, *x*_max_ the maximum *x* value for the spectrum and *x*_min_ the minimum *x* value for the spectrum. The normalised spectra had variable regions removed; these occur due to variable CO_2_ and H_2_O levels in the environment, (< 700 cm^-1^, between 2000 and 2500 cm^-1^ and > 3600 cm^-1^). The second derivative was taken, again using the Savitzky-Golay algorithm but, this time, with 41 neighbours and a 2^nd^ order polynomial. The preliminary smooth is not always needed, but if omitted, then more neighbours are required for optimum output during this step. It was observed that a less aggressive smoothing procedure at the beginning of the process, removed anomalous baseline corrections entirely.

### PCA

The normalised and corrected matrix of intensities underwent PCA using singular value decomposition with the *base* ***prcomp*** function in R.

During this process, the matrix was mean centred, but not scaled in any other way. Through comparison of the scree and loading plots, suitable PC scores were chosen to plot against each other as *x*y scatter graphs.

## Author contributions

M.A.S., E.A.Y., and A.D. designed the approach and interpreted the results. M.A.S, E.A.Y and A.D. defined the method, and A.D implemented it. M.A.S., E.A.Y., M.G. and A.D. analysed the data. M.G. provided glycosaminoglycan samples for analysis and M.G. and L.M. obtained relevant NMR and SAX-HPLC data. M.A.S., E.A.Y and M.G. supervised the study. All authors drafted and approved the manuscript.

## References

1. Casu, B., Naggi, A. & Torri, G. Re-visiting the structure of heparin. Carbohydr. Res. 403, 60–68 (2015).

2. Keire, D. A. et al. Diversifying the global heparin supply chain? Biopharm Int. 39, 2–8 (2015).

3. Casu, B. Structure and Biological Activity of Heparin. Adv. Carbohydr. Chem. Biochem. 43, 51–134 (1985).

4. Rabenstein, D. L. Heparin and heparan sulfate: Structure and function. Nat. Prod. Rep. 19, 312–331 (2002).

5. Casu, B. Structure and Active Domains of heparin. in Chemistry and Biology of Heparin and Heparan Sulfate 1–28 (2005). doi:10.1515/9783110289039.545

6. Casu, B. et al. Characterization of sulfation patterns of beef and pig mucosal heparins by nuclear magnetic resonance spectroscopy. Arzneimittelforschung. 46, 472–7 (1996).

7. Mauri, L. et al. Combining NMR Spectroscopy and Chemometrics to Monitor Structural Features of Crude Heparin. Molecules 22, (2017).

8. Liu, H., Zhang, Z. & Linhardt, R. J. Lessons learned from the contamination of heparin. Nat. Prod. Rep. 26, 313–321 (2009).

9. Guerrini, M. et al. Oversulfated chondroitin sulfate is a contaminant in heparin associated with adverse clinical events. Nat. Biotechnol. 26, 669–675 (2008).

10. Alban, S. et al. Comparison of established and novel purity tests for the quality control of heparin by means of a set of 177 heparin samples. Anal. Bioanal. Chem. 399, 605–620 (2011).

11. Keire, D. A., Mans, D. J., Ye, H., Kolinski, R. E. & Buhse, L. F. Assay of possible economically motivated additives or native impurities levels in heparin by 1H NMR, SAX-HPLC, and anticoagulation time approaches. J. Pharm. Biomed. Anal. 52, 656–664 (2010).

12. Fasciano, J. M. & Danielson, N. D. Ion chromatography for the separation of heparin and structurally related glycoaminoglycans: A review. J. Sep. Sci. 39, 1118–1129 (2016).

13. Hashii, N. et al. Heparin identification test and purity test for OSCS in heparin sodium and heparin calcium by weak anion-exchange high-performance liquid chromatography. Biologicals 38, 539–543 (2010).

14. Stanley, F. E. & Stalcup, A. M. The use of circular dichroism as a simple heparin-screening strategy. Anal. Bioanal. Chem. 399, 701–706 (2011).

15. Devlin, A., Mycroft-West, C., Guerrini, M., Yates, E. & Skidmore, M. Analysis of solid-state heparin samples by ATR-FTIR spectroscopy. bioRxiv 538074 (2019). doi:10.1101/538074

16. Sommers, C. D., Mans, D. J., Mecker, L. C. & Keire, D. A. Sensitive detection of oversulfated chondroitin sulfate in heparin sodium or crude heparin with a colorimetric microplate based assay. Anal. Chem. 83, 3422–3430 (2011).

17. Guerrini, M. et al. Orthogonal analytical approaches to detect potential contaminants in heparin. Proc. Natl. Acad. Sci. 106, 16956–16961 (2009).

18. Rudd, T. R. et al. Construction and use of a library of bona fide heparins employing 1H NMR and multivariate analysis. Analyst 136, 1380–1389 (2011).

19. Rudd, T. R. et al. How to find a needle (or anything else) in a haystack: Two-dimensional correlation spectroscopy-filtering with iterative random sampling applied to pharmaceutical heparin. Anal. Chem. 84, 6841–6847 (2012).

20. Tipson, R. S. Infrared Spectroscopy Of Carbohydrates A Review. National Bureau of Standards Monograph 110 (1968).

21. Grant, D., Long, W. F., Moffat, C. F. & Williamson, F. B. Infrared spectroscopy of chemically modified heparins. Biochem. J. 261, 1035–1038 (1989).

22. Harws, M. J. & Turvey, J. R. Sulfates of monosaccharides and derivatives Part VIII. Infrared spectra and optical rotations of glycoside sulfates. Carbohydr. Res. 15, 51–56 (1970).

23. Grant, D., Long, W. F., Moffat, C. F. & Williamson, F. B. Infrared spectroscopy of heparins suggests that the region 750-950 cm −1 is sensitive to changes in iduronate residue ring conformation. Biochem. J. 275, 193–197 (1991).

24. Lloyd, A. G., Dodgson, K. S., Price, R. G. & Rose, F. A. Infrared studies on sulfate esters I. Polysaccharide sulfates. Biochim. Biophys. Acta 46, 108–115 (1961).

25. Vasko, P. D., Blackwell, J. & Koenig, J. L. Infrared and raman spectroscopy of carbohydrates. Part I: Identification of OH and CH-related vibrational modes for D-glucose, maltose, cellobiose, and dextran by deuterium-substitution methods. Carbohydr. Res. 19, 297–310 (1971).

26. Grant, D., Moffat, C. F., Long, W. F. & Williamson, F. B. N Acetyl amide-II and methyl C—H bending infrared absorbances of chemically modified heparins. Biochem. Soc. Trans. 17, 502–503 (1988).

27. Neely, B. W. Infrared Spectra of Carbohydrates. Adv. Carbohyd. Chem. 12, 13–33 (1957).

28. Cabassi, F., Casu, B. & Perlin, A. S. Infrared absorption and raman scattering of sulfate groups of heparin and related glycosaminoglycans in aqueous solution. Carbohydr. Res. 63, 1–11 (1978).

29. Andrianova, V. M. & Zhbankov, R. G. Conformational analysis of carbohydrate nitrasted and the dependence of their vibrational spectrum on the type of nitro group rotamer. J. Struct. Chem. 28, 194–200 (1987).

30. Wiercigroch, E. et al. Raman and infrared spectroscopy of carbohydrates: A review. Spectrochim. Acta - Part A Mol. Biomol. Spectrosc. 185, 317–335 (2017).

31. Orr, S. F. D. Infra-Red Spectroscopic Studies of Some Polysaccharides. Biochim. Biophys. Acta 14, 173–181 (1954).

32. Grant, D., Moffat, C. F., Long, W. F. & Williamson, F. B. N-Sulphonate infrared absorbances of chemically modified heparins. Biochem. s 17, 498–500 (1989).

33. Mainreck, N. et al. Rapid Characterization of Glycosaminoglycans Using a Combined Approach by Infrared and Raman Microspectroscopies. J. Pharm. Sci. 100, 441–450 (2010).

34. Han, W., Li, Q., Lv, Y., Wang, Q. C. & Zhao, X. Preparation and structural characterization of regioselective 4-O/6-O-desulfated chondroitin sulfate. Carbohydr. Res. 460, 8–13 (2018).

35. Foot, M. & Mulholland, M. Classification of chondroitin sulfate A, chondroitin sulfate C, glucosamine hydrochloride and glucosamine 6 sulfate using chemometric techniques. J. Pharm. Biomed. Anal. 38, 397–407 (2005).

36. Grant, D., Long, W. F. & Williamson, P. B. A model of two conformational forms of heparins/heparans suggested by infrared spectroscopy. Med. Hypotheses 24, 131–136 (1987).

37. Grant, D., Moffat, C. F., Long, W. F. & Williamson, F. B. Carboxylate symmetric stretching infrared absorbances of chemically modified heparins. Biochem. Soc. Trans. 17, 500–501 (1988).

38. Casu, B., Scovena, G., Cifonelli, A. & Perlin, A. S. Infrared sprectra of glycosaminoglycans in deuterium oxide and deuterium chloride solution: quantiative evaluation of uronic acid and acetamidodeoxyhexose moieties. Carbohydr. Res. 63, 13–27 (1978).

39. Myron, P., Siddiquee, S. & Azad, S. Al. Partial structural studies of fucosylated chondroitin sulfate (FuCS) using attenuated total reflection fourier transform infrared spectroscopy (ATR-FTIR) and chemometrics. Vibrational Spectroscopy 89, (Elsevier B.V., 2017).

40. Vasconcelos Oliveira, A. P. et al. Characteristics of Chondroitin Sulfate Extracted of Tilapia (Oreochromis niloticus) Processing. Procedia Eng. 200, 193–199 (2017).

41. Rudd, T. R. et al. Unravelling structural information from complex mixtures utilizing correlation spectroscopy applied to HSQC spectra. Anal. Chem. 85, 7487–7493 (2013).

